# MHC-II expressing neutrophils circulate in blood and milk during mastitis and show high microbicidal activity

**DOI:** 10.1101/2022.09.01.506187

**Authors:** Marion Rambault, Florence B. Gilbert, Philippe Roussel, Alexia Tessier, Valérie David, Pierre Germon, Nathalie Winter, Aude Remot

## Abstract

Bovine mastitis are mainly caused by bacterial infection. They are responsible for economic losses and have an impact on the health and welfare of animals. The increase in the somatic cell count in milk during mastitis is mainly due to the influx of neutrophils which have a crucial role in the elimination of pathogens. For a long time, these first line defenders has been view as microbes’ killers with limited role in the orchestration of the immune response. However, their role is more complex and we recently characterized a MHC-II expressing neutrophil subset with regulatory capacities in cattle. In this study, we questioned the implication of different neutrophils subsets in the mammary gland immunity during clinical and subclinical mastitis. Here, we described for the first time that, in blood as in milk, neutrophils are a heterogeneous population and encompass at least two subsets distinguishable with their expression of MHC-II. We observed higher bactericidal capacities of milk MHC-II^pos^ neutrophils as compared to their classical counterparts, due to a higher production of ROS and phagocytosis ability. MHC-II^pos^ neutrophils are enriched in milk during a subclinical mastitis as compared to blood. Moreover, we observed a positive and highly significant correlation between MHC-II^pos^ neutrophils and T lymphocytes present in milk during subclinical mastitis. To conclude, our study could open the way to the discovery of new biomarkers of mastitis inflammation.

## Introduction

Mastitis is an inflammation of the mammary gland resulting mostly from bacterial infection. It affects the health and welfare of animals and leads to economic losses due to lower milk production and quality, antibiotic treatments and premature drying off and culling (Halasa, Huijps et al. 2007). Mastitis, the most prevalent and costly disease in dairy herds worldwide, presents a spectrum of clinical conditions ranging from severe clinical mastitis to subclinical mastitis where the animals do not show signs of discomfort. When clinical mastitis declares, local signs of mammary inflammation such as swollen and red mammary gland, with painful and warm udder which are a concern for the welfare of the animal, is accompanied by lumps in milk which renders it unproper to consummation. By contrast, subclinical mastitis displays no apparent clinical signs of inflammation while milk somatic cell count (SCC) is increased to levels higher than 200,000 cells/mL which impacts milk quality and lowers the price (Smith, Hillerton et al. 2001). Therefore, subclinical mastitis represents 70% to 80% of economic losses linked to mastitis (Goncalves, Lyman et al. 2017) and measures to better understand and prevent these forms of the diseases are also needed.

The SCC that provides an indication of the inflammatory status of the mammary gland is now a well-established parameter in milk control. Numbers above 150,000 - 200,000 cells/mL milk are often set as a marker to identify mastitis, even though this threshold varies depending on countries (Smith, Hillerton et al. 2001). The increase of the SCC during mastitis is mainly due to the influx of polymorphonuclear neutrophils (PMNs) from blood to the mammary gland (Rainard and Riollet 2003, Koess and Hamann 2008, Schwarz, Diesterbeck et al. 2011). PMNs are rapidly recruited during the acute inflammatory response to defend the mammary gland against invading pathogens and represent the signature of inflammation by being the predominant cell type in milk (Paape, Wergin et al. 1979, Rainard and Riollet 2003). In addition to PMNs, macrophages and lymphocytes are also present in milk and taken into account in the SCC. PMNs to macrophages and lymphocytes ratio is positively correlated to inflammation, macrophages being considered as the major leucocyte population in healthy milk (Rambault, Roussel et al. 2021).

Since decades, genetic tools are available to help dairy farms to control mastitis. Somatic cell score (SCS, a logarithmic transformation of SCC) has been used to select mastitis resistant cows (Rupp, Bergonier et al. 2009, Brand, Hartmann et al. 2011, Weigel and Shook 2018). A divergent selection experiment in goats based on somatic cell score (SCS) showed favorable results with reduction of clinical and subclinical mastitis in the low SCS group and higher incidence in the high SCS group (Rupp, Bergonier et al. 2009). Other authors conducted a divergent genetic selection on Holstein and Normande breeds based on sire breeding values. In the Holstein breed, a lower SCS and a lower prevalence of clinical mastitis in the mastitis resistant group was observed as compared to the control group. However, results in the Normande breed were much less marked in first lactation (Lefebvre, Barbey et al. 2020).

While divergent selection based on genetic or SCS seems promising to select mastitis resistant cows, improvements in the selection criteria are still required, especially to refine cell phenotyping, and more particularly PMNs’, when measuring SSC in different breeds.

PMNs are efficient phagocytes and release a large amount of reactive oxygen species (ROS) into phagocytic vacuoles to rapidly kill pathogens (Nguyen, Green et al. 2017). Generally described as an innate cell type with pro-inflammatory roles, some studies in humans and mouse models, highlighted new phenotypes and functions for unconventional PMNs, such as suppression of T cells’ proliferation (Mantovani, Cassatella et al. 2011, Pillay, Tak et al. 2013). In cattle, we recently characterized a new subset of PMNs expressing the major histocompatibility complex class II (MHC-II) molecules. These unconventional MHC-II^pos^ PMNs were identified in bone marrow, spleen and blood of healthy cows. Classical MHC-II^neg^ PMNs and MHC-II^pos^ PMNs displayed distinct transcriptomic profiles and functions. Importantly, we also showed that MHC-II^pos^ PMNs were able to inhibit the proliferation of T cells *in vitro* whereas classical PMNs were not. Because of this function, we proposed to name them bovine regulatory PMNs (Rambault, Doz-Deblauwe et al. 2021).

The presence of regulatory PMNs during infection remains to be investigated. Therefore, we decided to address this question in mastitis. We followed both Normande and Holstein cows during several lactations where different episodes of clinical or subclinical mastitis occurred. We analyzed the two MHC-II^neg^ and MHC-II^pos^ PMNs subsets in blood and milk. We demonstrated that both subsets were present in milk during mastitis, and we observed higher bactericidal capacities of milk MHC-II^pos^ PMNs as compared to their classical counterparts. Interestingly, we also observed a positive correlation between counts of MHC-II^pos^ PMNs and T cells in milk during subclinical mastitis.

## Materials and Methods

### Studied herds and animal protocols

Three French dairy herds were recruited for this study. They are located in Centre-Val de Loire and Normandie areas. All herds hosted cows from the Holstein breed and one also hosted cows from the Normande breed. For all herds, procedures were approved by local Ethics Committee (agreements APAFIS#29498-2021020410061759 and APAFIS#23901-20200203095344). Our protocol did not interfere with farms breeding practices. All animals used in our study remained in the herd with their fellows, without any space or food restriction.

#### (A) Blood and milk sampling from healthy Holstein and Normande cows

Twelve healthy Holstein and Normande cows (see Figure 1) were all in early second lactation (60-94 days after calving). Milk was aseptically collected and analyzed to confirm the absence of inflammation in the mammary gland. Blood sampling was performed with 10 ml vacutainer K2 EDTA tubes for flow cytometry and 4 ml vacutainer K3 EDTA tubes for complete blood count. Thirty ml of milk were collected from each quarter of Holstein cows with no clinical signs of mastitis.

**Figure 1:**
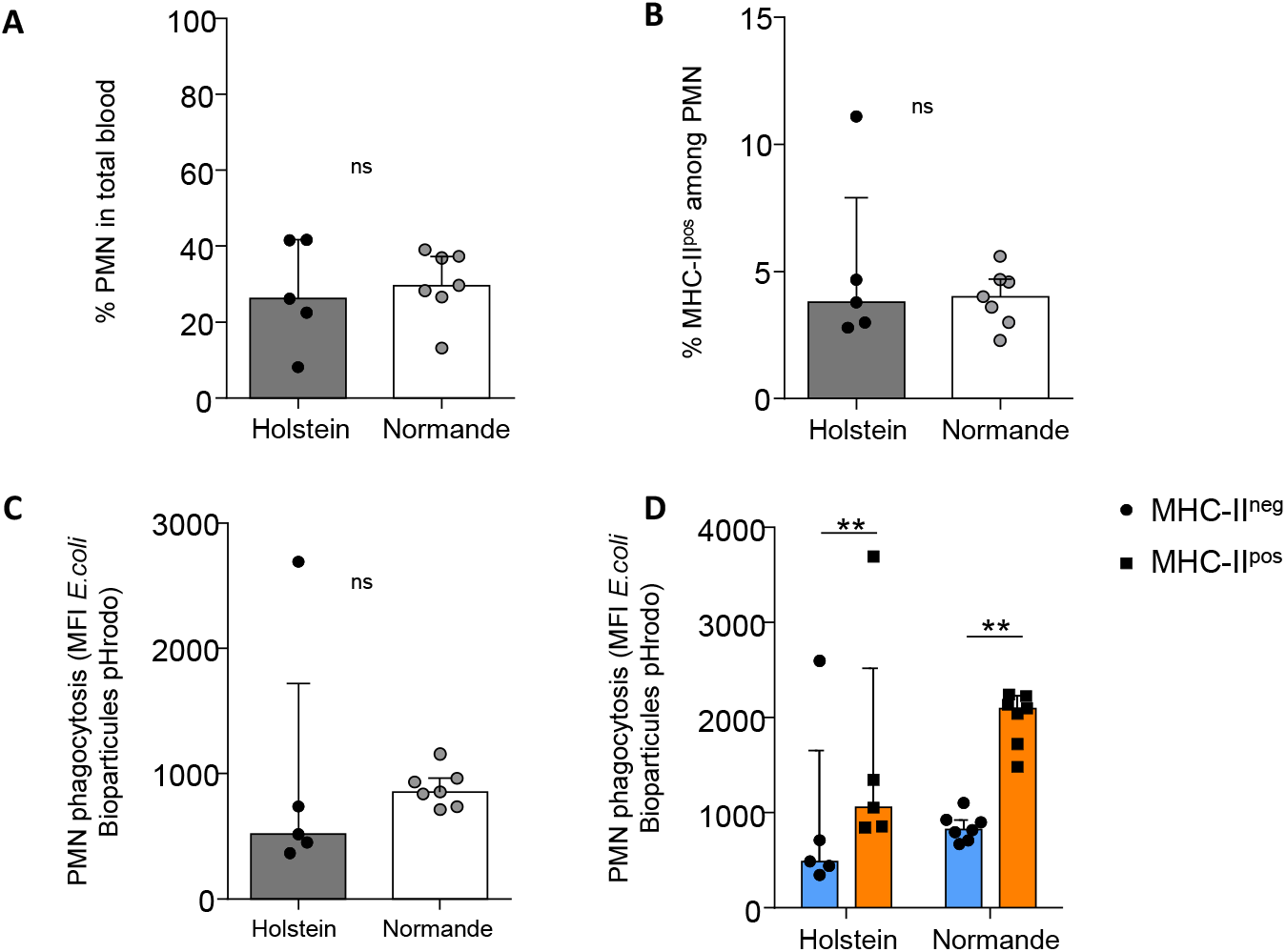
Blood MHC-II^pos^ PMN display better phagocytosis activity as compared to MHC-II^neg^ PMN in Holstein and Normande cows. Cells were prepared from blood from the two breeds **(A)** Proportion of total PMNs in blood was determined by complete blood count and **(B)** percentage of MHC-II^pos^ PMN was determined by flow cytometry among CD45+ G1+ live cells **(C, D) P**hagocytosis of pHrodo *E. coli* bioparticles by total PMNs **(C)** or MHC-II^pos^ and MHC-II^neg^ PMNs was assessed by measuring fluorescence intensity directly correlated to ingested particles. (**A-D**) Data are median with interquartile range, n = 5 Holstein, n = 7 Normande; data are representative of 1 experiment over 2 independent. \*\**P*<0.01; \*\*\**P*<0.001; ns, not significant (Wilcoxon signed-rank test).

#### (B) Collection of samples from cows with clinical mastitis

As soon as milking technicians identified a cow with a clinical mastitis, the animal was recruited for the experiment and milk from the infected quarter and blood were collected at different time points, as described in **Figure 3A**. After the diagnosis of clinical mastitis and the first milk sampling, MASTIJET® (MSD) was administered intramammarily during 4 consecutive milkings. A total of ten cows diagnosed with clinical mastitis were followed.

#### (C)Collection of samples from cows with subclinical mastitis

Milk of each quarter and blood were collected for 20 lactating cows suspected of having subclinical mastitis based on composite milk SCC above 300,000 cells/mL at the last individual cow milk cell count and with a normal milk aspect the week before. Quarters with SCC above 150,000 cells/mL and with a bacterial culture positive were considered to have a subclinical mastitis.

### Cell preparation from blood samples

Vacutainer K2 EDTA tubes (10 ml) were centrifuged at 1,000g for 10 min at 20°C. Plasma and buffy coat were discarded, then ACK (Gibco) (5 vol/1 vol of blood cell) was added for 5 min at room temperature to lyse red blood cells. Cells were washed twice in D-PBS (without calcium and magnesium) with 2mM EDTA.

### Milk samples

After cleaning and disinfection of the teat, the first 2 streams of milk were discarded and 30 ml of milk of each quarter were collected in sterile tubes. For animals with mastitis, milk was collected before any antibiotic treatment.

#### Microbial culture and bacterial identification

Thirty microliters of milk from all quarters were plated on blood agar plate (Oxoid). Plates were incubated at 37°C and examined at 24h and 48h. Bacterial identification was performed using conventional bacteriology technics as described in Laboratory Handbook on Bovine Mastitis, 3rd Edition (Adkins, Middleton et al. 2017). If conventional bacteriology tests were not sufficient for bacterial identification, MALDI-TOF mass spectrometry was performed. If three or more types of colonies were visible after incubation, milk sample was considered as contaminated and no identification was performed.

#### Preparation of cells from milk samples

Milk somatic cell count (SCC) was determined with an automated cell counter (Fossomatic model 90, Foss Food Technology, Hillerod, Denmark). Milk was diluted two times in D-PBS with 2% (v/v) of horse normal serum (Gibco) and centrifuged at 1,000g for 10 min at 4°C. The cream layer and supernatant were discarded and the cell pellet was resuspended and washed twice in D-PBS with 2% of horse normal serum (400g, 10 min, 4°C).

### Flow cytometry

Cells prepared from the milk and blood samples were suspended in D-PBS (wo Ca/Mg) containing 10% (v/v) of horse normal serum (Gibco) and 2mM EDTA, and labeled for 30 min with primary antibodies (30 min, 4°C). After washes in D-PBS with 10% of horse normal serum (400g, 10 min, 4°C), cells were labeled with the corresponding fluorescent-conjugated secondary antibodies. Two panels of antibodies were designed for the analysis of markers of myeloid or lymphoid cells. All antibodies used are listed in **Table 1**. Cells were washed and fixed with BD cell fix solution diluted four times in D-PBS (wo Ca, Mg). Data were acquired with a LSR Fortessa™ X-20 Flow cytometer (Becton Dickinson) and results analyzed with Kaluza software (Beckman Coulter).

**Table 1:**
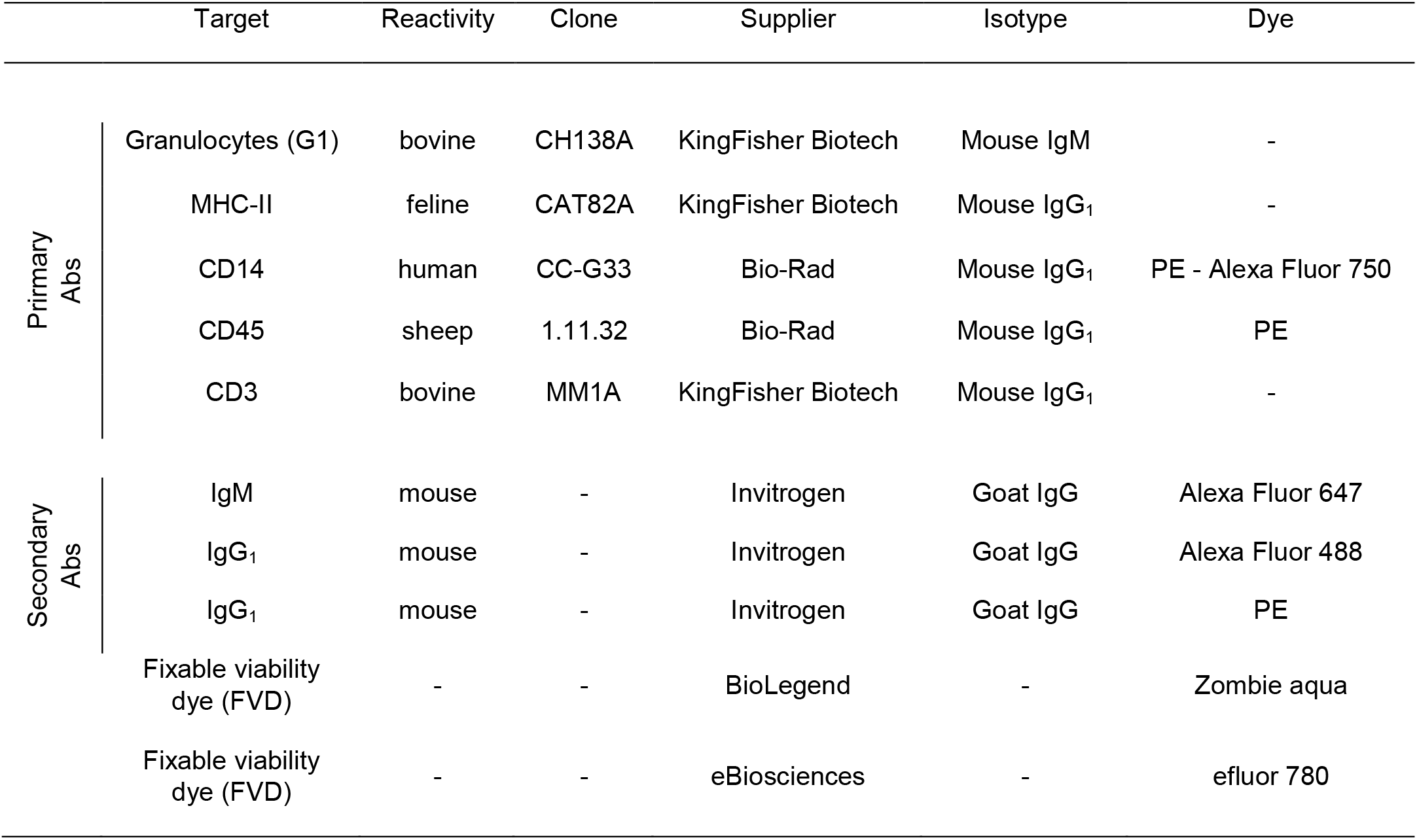
Antibodies used for flow cytometry analysis.

### Phagocytosis

Following the labelling of the cells with the antibodies recorded in Table 1, two different methods were used to assess the phagocytosis capacity of PMNs.

#### pHrodo Red E.coli BioParticules

Phagocytosis was measured using pHrodo™ Red *E. coli* BioParticles® Conjugate (Molecular Probes®) following the manufacturer’s instructions. Briefly, 10^5^ cells/well were incubated in a 96 wells microplate for 1 h at 37°C with 20 µg/well of pHrodo *E. coli* BioParticles in RPMI-1640 supplemented with 2 mM L-glutamine, 10 mM HEPES, and 1mg/ml of BSA with extremely low endotoxin level (≤1.0 EU/mg) (hereafter referred to as RPMI complete medium). The fluorescence linked to the ingestion of the *E. coli* BioParticles was directly measured with the LSR Fortessa™ X-20 Flow cytometer.

#### GFP-E. coli P4

*Escherichia coli* strain P4 was transformed with the *gfp*-carrying plasmid pFPV25.1 (Bramley 1976, Valdivia and Falkow 1996). Transformants were grown in BHI medium in the presence of 100µg/ml ampicillin. Then, we used *E*.*coli* P4-GFP strain to test the phagocytosis capacity of PMNs. *E*.*coli* P4-GFP strain were grown in 10 ml BHI medium with 100µg/ml ampicillin overnight at 37°C without agitation. Bacteria were then diluted in BHI medium (1 vol/100 vol) and incubated for 6 h at 37°C without agitation. Bacterial concentration was determined by the optical density at 600 nm and adjusted at 8.10^6^ CFU/ml in RPMI complete medium. After milk cells labelling, 2.10^5^ of cells were plated in a 96 wells microplate and infected at a MOI of 2 in RPMI complete medium for 30 min at 37°C or at 4°C.

### Reactive oxygen species (ROS) production

ROS produced by PMNs were quantified using the CellROX® Orange Flow Cytometry Assay Kits (MolecularProbes®, C10493) following the manufacturer’s instructions. Briefly, 10^5^ milk cells were first incubated for 1 h at 37°C in RPMI complete medium with or without 400 µM of TBHP in a 96-wells black microplate and then for 30 min with 100 nM CellROX® and 1µL/mL of efluor780 fixable viability dye. Fluorescence was directly measured with the LSR Fortessa™ X-20 Flow Cytometer.

### Statistical analysis

Individual data and the median were presented in the figures. Statistical analyses were performed with R Studio and Prism 6.0 software (GraphPad). Wilcoxon signed-rank test and Spearman correlation test were used. Represented p-values were: \**P* < 0.05; \*\**P* < 0.01; and \*\*\**P* < 0.001; \*\*\*\**P* < 0.0001.

## Results

### MHC-II^pos^ PMNs with high phagocytic capacity and ROS production similarly circulate in blood of healthy Holstein and Normande cows

In order to better characterize PMNs subsets in dairy cows, we analyzed their proportions in the blood of healthy Holstein and Normande cows, the two main dairy breeds in western France. To limit physiological variations due to different stages of lactation, we selected cows within the first trimester of the second lactation. We then analyzed MHC-II^pos^ and MHC-II^neg^ PMNs according to the gating strategy as described in our previous study (Rambault, Doz-Deblauwe et al. 2021). We did not see any difference between the two breeds for the ratio of total PMNs circulating in blood (**Figure 1A**). Similarly, the ratio of MHC-II^pos^ among total PMNs was comparable for the two breeds (**Figure 1B**).

We next compared by flow cytometry the phagocytic activity of PMNs from Holstein and Normande cows using conjugated pHrodo™ Red *E. coli* BioParticles (Rambault, Doz-Deblauwe et al. 2021) allowing us to quantify phagocytosis by mean fluorescence intensity (MFI) as a direct correlation of ingested particles. Phagocytosis by total PMNs was comparable for both breeds (**Figure 1C**). Interestingly, for both breeds, MHC-II^pos^ PMNs were significantly more efficient phagocytes as compared to MHC-II^neg^ PMNs (**Figure 1D**).

### MHC-II^pos^ PMNs with high microbicidal activities are recruited to the mammary gland

It is established that PMNs represent the largest proportion in SSC during mastitis. Yet, the presence of different subsets of PMNs in SSC has not been investigated. Because we did not record difference between breeds, we concentrated our analysis of milk SCC on Holstein cows, the most represented breed in French dairy herds. We analyzed the presence of MHC-II^neg^ and MHC-II^pos^ PMNs in milk according to the gating strategy described in **Figure 2A**. As in blood, the MHC-II^pos^ subset of PMNs was also observed in milk. Next, we compared phagocytosis by the two subsets of milk PMNs using an *E. coli* P4 strain expressing the fluorescent GFP protein. After incubation of milk cells with *E*.*coli* P4-GFP at 37°C, we analyzed by flow cytometry the percentage of the two subsets of PMNs carrying green fluorescence. We normalized this ratio on cells stimulated with *E*.*coli* P4-GFP at 4°C where phagocytosis is blocked (Peterson P.K., Verhoef J et al. 1977). As in blood, milk MHC-II^pos^ PMNs significantly phagocytosed more bacteria as compared to MHC-II^neg^ PMNs (*P*=0.0156) (**Figure 2B**). As ROS production is an important mechanism used by PMNs to kill bacteria, we also measured the potential of milk cells to produce these bactericidal molecules after incubation with the nonspecific chemical inducer *tert-butyl hydroperoxide* (TBHP). Milk MHC-II^pos^ PMNs produced significantly more ROS than MHC-II^neg^ PMNs (*P*=0.0078) (**Figure 2C**). Therefore, as observed in blood, MHC-II^pos^ PMNs recruited to the mammary gland were better equipped with bactericidal arsenal as compared to their classical MHC-II^neg^ counterparts.

**Figure 2:**
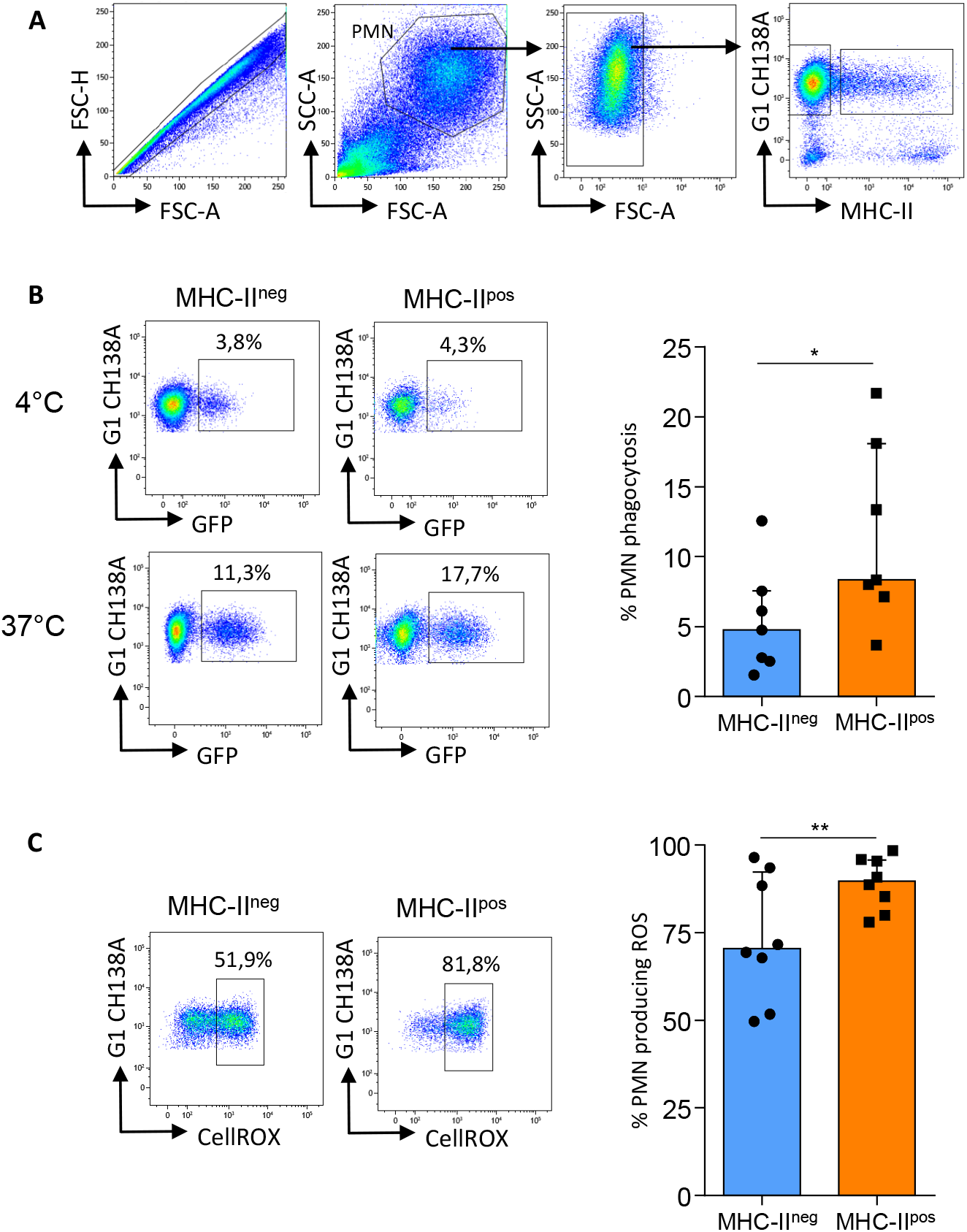
MHC-II^pos^ PMN have a higher bactericidal capacities compare to MHC-II^neg^ PMN. **(A)** Flow cytometry gating strategy used to identify MHC-II^neg^ and MHC-II^pos^ PMNs in milk. **(B)** After milk cells labelling, phagocytosis by or MHC-II^pos^ or MHC-II^neg^ PMN was assessed using *E. coli* P4-GFP strain at 37°C. PMNs phagocytosis were normalized on PMNs stimulated with *E*.*coli* P4-GFP at 4°C. Data represent n=2 experiments with n=7 different cows. **(C)** Oxidative stress was measured in MHC-II^pos^ and MHC-II^neg^ neutrophils using the CellROX Orange probe that reacts with all ROS species. Cells were activated with TBHP and levels of ROS were measured by flow cytometry. Data represent n=2 experiments with n=8 different cows. **(B, C)** Fluorescence for each sample is depicted, and paired MHC-II^pos^ and MHC-II^neg^ samples were analyzed for each animal. Data are median with interquartile range. \**P*<0.05; \*\**P*<0.01 (Wilcoxon signed-rank test)

### Ratios of MHC-II^pos^ and MHC-II^neg^ PMNs circulating in blood and recruited in milk do not vary during the course of a clinical mastitis episode

Since total PMNs in SCC are a marker of mastitis progression after treatment with antibiotics, we next addressed how MHC-II^neg^ and MHC-II^pos^ subsets behaved during a clinical mastitis episode. In order to do that, we followed 10 cows as soon as a clinical mastitis episode was detected. According to animal care protocols, antibiotic treatment was administered into the mammary gland as soon as the episode was detected. We sampled blood and milk from the infected quarter at three different time points: S1 (day 0), S2 (between day 3 and 6) and S3 (day 21) (**Figure 3A**). Most clinical mastitis episodes were associated to mild symptoms and only Gram-positive bacteria were detected after microbial culture (**Table 2**). Two bacterial identifications were unsuccessful due to milk contamination and no bacteria were detected in one milk sample. As generally observed for clinical mastitis, inflammation signs at S1 were high and SCC ranged from 1,000 × 10^3^ to 35,000 × 10^3^ cells/mL. At S2, from 3 to 6 days after antibiotic treatment, milk SCC was generally lower: 4 quarters had less than 300,000 cells/mL, from which 3 displayed negative microbial culture. At S3 (day 21), milk SCC from 6 quarters was inferior to 300,000 cells/mL and displayed negative microbial culture indicating recovery. When bacteria were detected in S2 and/or S3 milk samples, they belonged to the same species as isolated from S1; therefore, we considered that bacteria resisted antibiotic treatment (4/10 cows).

**Figure 3:**
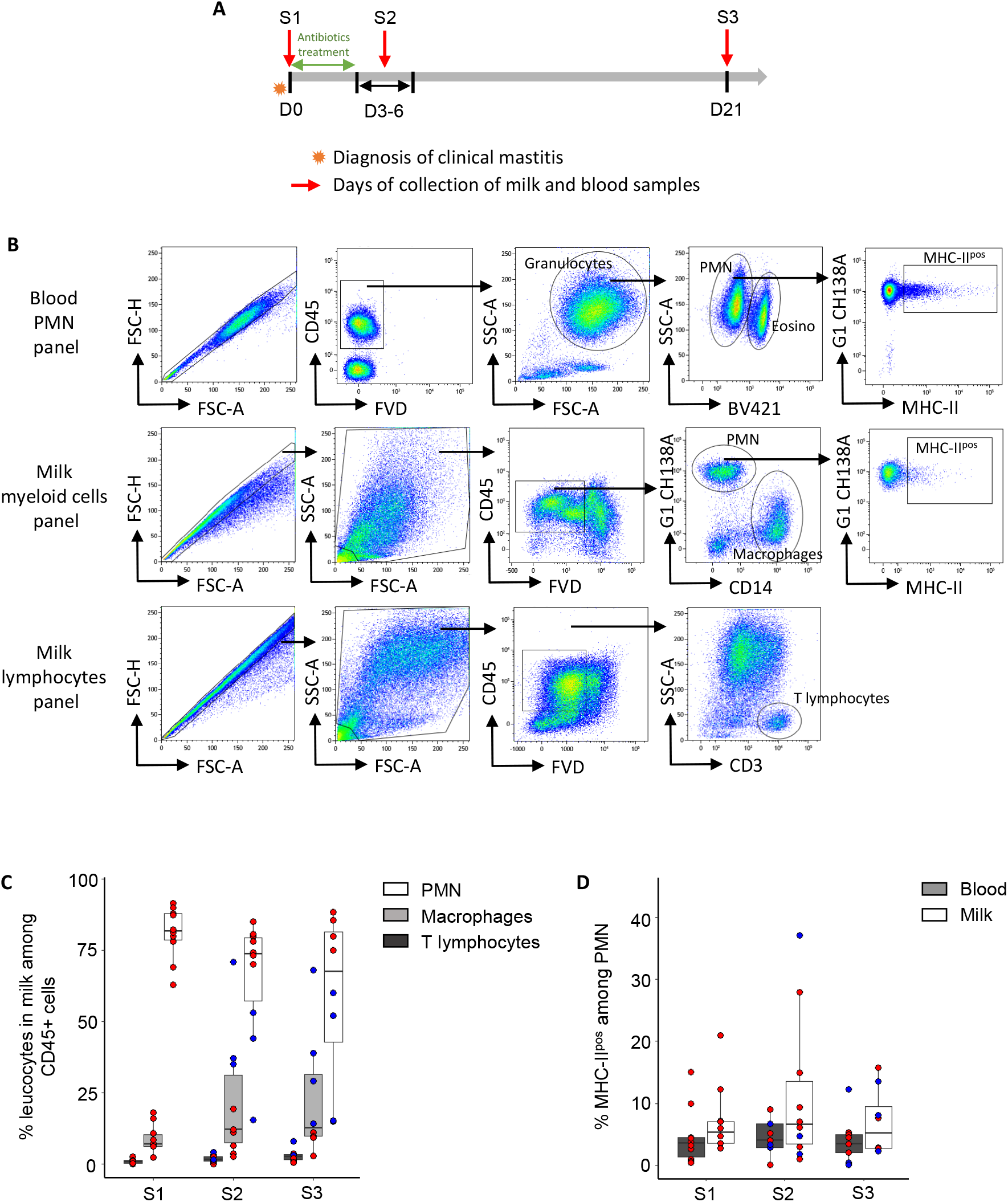
MHC-II^pos^ PMN are present in blood and milk during clinical mastitis and their proportion does not vary over time. **(A)** Description of sampling protocol. A first blood sample and milk sample from the infected quarter (S1) were immediately collected after the diagnosis of the clinical mastitis. After an antibiotic treatment for 5 milking of the infected quarter, a second blood and milk samples (S2) were collected between day 3 and day 6 (the day was choose to avoid week-ends). Finally, a third blood and milk samples were collected at day 21. **(B)** Flow cytometry gating strategy used to identify different cells populations in blood and in milk. Blood MHC-II^pos^ PMN are defined as CD45+ SCC_high_ FSC_high_ G1+ MHC-II+. Eosinophils were excluded by excitation with violet laser in the “granulocytes” gate. Milk PMN, macrophages, and T lymphocytes are defined as CD45+ G1+ CD14-, CD45+ G1-CD14+ and CD45+ CD3+ respectively. Milk MHC-II^pos^ PMN are defined as MHC-II+ in the “PMN” gate. **(C)** Percentage of PMN, macrophages and T lymphocytes in milk at each sample collection time. Red dots = milks with SCC > 300 000 cells/mL; bue dots = milks with SCC < 300 000 cells/mL. **(D)** Percentage of MHC-II^pos^ PMN in blood and in milk at each time point. **(C, D)** Data are boxes and whiskers min to max. n = 10 cows and 27 independent experiments.

**Table 2:** Description of clinical mastitis cases. Lactation stage: Near calving (< 7 days after calving), Early (7 < days after calving < 100), Mid (100 < days after calving < 200), Late (> 200 days after calving) Clinical grade: 1 = slight damage (lumps in milk only); 2 = moderate damage (swollen and painful udder) Microbial culture: + = presence of colonies on blood agar, - = no colonies on blood agar

| Clinical mastitis (CM) | Lactation stage | Lactation number | Pathogen | Clinical grade | S1 |  | S2 |  | S3 |  |
| --- | --- | --- | --- | --- | --- | --- | --- | --- | --- | --- |
|  |  |  |  |  | SCC x 10<sup>3</sup> | Microbial culture | SCC x 10<sup>3</sup> | Microbial culture | SCC x 10<sup>3</sup> | Microbial culture |
| CM1 | Near calving | 7 | *S.aureus* | 1 | 8400 | + | 10500 | - | 1900 | - |
| CM2 | Early | 7 | *Enterococcus spp* | 1 | 8000 | + | 632 | - | 83 | - |
| CM3 | End | 2 | *S.uberis* | 1 | 4800 | + | 1500 | + | 14600 | + |
| CM4 | End | 1 | *Staphylococcus arletae* | 1 | 3400 | + | 6000 | - | 13200 | + |
| CM5 | Early | 5 | Contaminated sample | 1 | 6400 | + | 63 | - | 7 | - |
| CM6 | Mid | 3 | *S.aureus* | 2 | 35000 | + | 31000 | + | 24000 | + |
| CM7 | End | 7 | No bacteria | 2 | 9000 | - | 257 | - | 81 | - |
| CM8 | Near calving | 1 | *S.uberis* | 1 | 26500 | + | 6300 | + | 78 | - |
| CM9 | End | 4 | Contaminated sample | 1 | 1000 | + | 316 | - | 41 | - |
| CM10 | Early | 1 | *S.uberis* | 1 | 5200 | + | 78 | - | 45 | - |

Next, we thoroughly analyzed blood and milk SCC composition by flow cytometry and designed three antibodies labeling panels and gating strategies to distinguish key cells involved in mastitis physiopathology, including the two PMNs’ subsets (**Figure 3B**) that we followed in blood and milk. The myeloid panel allowed us to distinguish the two subsets of PMNs from macrophages and the lymphoid panel informed us on CD3 lymphocytes. We could not perform flow cytometry analysis on two S3 milk samples because of too low SCC. As expected, in S1 milk samples, PMNs were the main cell type whereas low proportions of macrophages and T lymphocytes were detected (**Figure 3C**). In S2 and S3, due to start of recovery after treatment, ratios of the different cell types were more heterogeneous as compared to S1. We distinguished in our analysis samples with SCC superior to the threshold of 300,000 cells/ml indicating that the episode of clinical mastitis was still ongoing (**Figure 3C**, red dots) from samples with SCC inferior to the threshold as a sign of recovery (**Figure 3C**, blue dots). In S2 and S3, PMNs were still the main cell type for most samples. However, in three S2 milk samples and four S3 milk samples, PMNs ratios started to decrease which correlated with SCC below the threshold of 300,000 cells/ml. The samples with lowest SCC had also the highest proportion of macrophages, indicating that the mammary gland was returning to homeostasis. Interestingly, no difference in proportion of T lymphocytes was observed between the 3 time points. MHC-II^pos^ PMNs were present in blood and milk at each time point (**Figure 3D**). There was a trend towards more elevated MCH-II^pos^ to MHC-II^neg^ PMNs ratios in milk as compared to blood although this did not reach statistical significance due to high heterogeneity. We could not correlate the MCH-II^pos^ to MHC-II^neg^ PMNs ratios to the threshold of 300,000 cells/ml in SCC. Moreover, the proportion of MHC-II^pos^ PMNs did not seem to be related to bacterial clearance (see Table 2). In conclusion, even though we detected the new subset of MHC-II^pos^ regulatory neutrophils in blood and milk during each clinical mastitis episode, we could not correlate their presence to the level of inflammation, the sampling time (onset versus treated -mastitis) or bacterial clearance.

### MHC-II^pos^ regulatory PMNs positively correlate with T cells in milk during subclinical mastitis

Subclinical mastitis is characterized by absence of visible inflammation of the mammary gland. Yet, PMNs still account for most cells in the milk SCC. Thus, we investigated if MHC-II^pos^ PMNs were present in milk and blood during this mild form of mastitis. In order to do so, we selected 20 Holstein cows presenting a composite milk SCC above 300,000 cells/mL the week before our experiments based on information provided by monthly milk quality control. On the day of the experiment, milk samples of normal aspect were collected from individual quarters. After microbial culture and SCC, we defined subclinical mastitis in each quarter of the mammary gland based on positive microbial culture and SCC above 150,000 cells/mL on that day. Three cows showed no sign of subclinical mastitis and were excluded. Among the 17 remaining animals sampled (i.e. 68 quarters), 27 quarters were classified with subclinical mastitis with SCC ranging from 168 × 10^3^ to 28,858 × 10^3^ (**Table 3**). Proportions of different leucocytes present in blood and milk were analyzed by flow cytometry (see Figure 3B for gating strategy). As expected, total PMNs were still the main cell type among total milk leukocytes during subclinical mastitis while lower proportions of macrophages and T lymphocytes were observed (**Figure 4A**). Interestingly, among total PMNs, the MHC-II^pos^ subset was present in a significantly higher proportion in milk as compared to blood (**Figure 4B**, *P*=0.0042). Thus, contrary to what was observed during clinical mastitis, the regulatory PMN subset seemed to accumulate in milk during subclinical mastitis. Moreover, we observed a positive and highly significant correlation between counts of MHC-II^pos^ PMNs and T lymphocytes present in milk (**Figure 4C**, *P*<0.0001).

**Table 3:** Bacterial identification and somatic cell count of subclinical mastitis cases. LF = left front; RF = right front; LR = left rear; RR = right rear; CNS = coagulase negative staphylococci; Contaminated sample = three or more types of colonies.

| Subclinical mastitis (SM) | Quarter | SCC x 10<sup>3</sup> | Pathogen(s) |
| --- | --- | --- | --- |
| SM1 | LR | 4705 | CNS |
|  | LF | 423 | CNS |
| SM2 | RR | 699 | Contaminated sample |
| SM3 | LR | 842 | CNS |
|  | LF | 627 | CNS |
| SM4 | RR | 2703 | CNS |
|  | LR | 759 | CNS |
|  | RF | 1066 | CNS |
|  | LF | 677 | *Corynebacterium spp* |
| SM5 | LF | 981 | Yeast |
| SM6 | RR | 440 | CNS + *Enterococcus spp* |
|  | LF | 508 | CNS + *Corynebacterium spp* |
| SM7 | LF | 2769 | CNS |
| SM8 | LR | 9389 | *Streptococcus uberis* |
|  | LF | 311 | No bacteria |
| SM9 | RR | 2966 | CNS |
| SM10 | LR | 28858 | *Staphylococcus arlettae* |
| SM11 | LR | 169 | CNS |
|  | RR | 5576 | *S. aureus* |
| SM12 | RF | 203 | *S. aureus* |
| SM13 | LR | 3457 | *S. aureus* |
| SM14 | RF | 1738 | Contaminated sample |
| SM15 | LR | 2618 | No bacteria |
| SM16 | RF | 915 | *S. aureus* |
|  | LR | 361 | No bacteria |
| SM17 | LR | 606 | *S. aureus* |
|  | RR | 491 | *S. aureus* |

**Figure 4:**
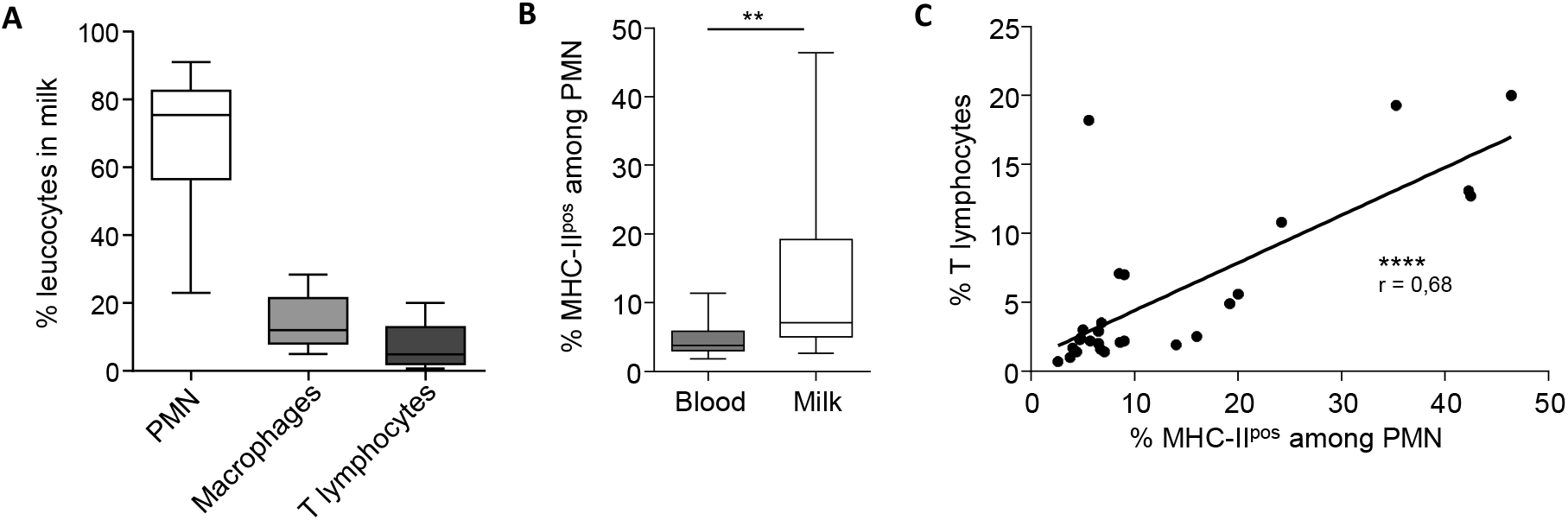
MHC-IIpos PMN accumulate in milk during subclinical mastitis and correlate with T cells. **(A)** Percentage of PMN, macrophages and T lymphocytes among CD45+ cells in milk during subclinical mastitis. **(B)** Percentage of MHC-II^pos^ PMN in blood and in milk during subclinical mastitis. **(A, B)** Data are boxes and whiskers min to max. **(C)** Correlation between the percentage of T lymphocytes and MHC-II^pos^ PMN in milk during subclinical mastitis. Data represent n = 2 independant experiments with n = 27 quarters from 17 cows. \*\**P*<0.01; \*\*\*\**P*<0.0001 (Wilcoxon signed-rank test) **(B)**, Spearman correlation test with 95% confidence interval **(C)**).

## Discussion

In agreement with a long-record of data on mastitis and follow up of the high milk SCC as a marker of mammary gland inflammation, we observed in our study that total PMNs represented the most abundant leucocytes in milk on the day of clinical mastitis diagnosis (Leitner, Shoshani et al. 2000, Schwarz, Diesterbeck et al. 2011), and also 21 days after, in most cases. However, we described here for the first time that, in blood as in milk, PMNs during mastitis are an heterogenous population and encompass at least two subsets. In humans and mice, PMNs heterogeneity is now well established (Silvestre-Roig, Hidalgo et al. 2016, Li, Wang et al. 2019). In cattle, despite PMNs being main cells recruited to the mammary gland during mastitis, they are still simply described as G1 positive cells (anti-granulocyte CH138A antibody) (Leitner, Shoshani et al. 2000, Leitner, Eligulashvily et al. 2003, Piepers, De Vliegher et al. 2009). We recently described in healthy cattle a new population of regulatory PMNs characterized by MHC-II expression and T-cell suppression capacity (Rambault, Doz-Deblauwe et al. 2021). Here, we show that this same population is present in milk during clinical and subclinical mastitis. Of note, the proportion of milk MHC-II^pos^ PMNs was highly variable in both forms of the disease and a possible explanation is the diversity of pathogens involved inducing different levels of inflammation between animals. On the same ground, it is worth pointing out that all clinical mastitis episodes were not cured at d21 as sometimes pathogens were still detected in milk with remaining high SCC. These heterogenous clinical situations might also explain our variable results on the PMN MHC-II^pos^ versus MHC-II^neg^ ratio. For future investigations, it would be interesting to include longer time points, to see if the MHC-II^pos^ subset increases in proportion when the acute phase of the mastitis episode is over. Indeed, during our study of clinical mastitis limited to 21 days, MHC-II^pos^ PMNs were observed at each sampling time point – at onset of the disease of after antibiotic treatment and beginning of recovery-in blood and milk and we could not correlate their presence with the course of the disease.

Among differences in functions between the two subsets, we demonstrated here that MHC-II^pos^ PMNs had higher phagocytic and ROS production capacities as compared to classical MHC-II^neg^ PMNs. In our previous report, we observed that blood MHC-II^pos^ PMNs were higher producers of ROS than their MHC-II^neg^ counterparts but we could not observe any difference in phagocytosis (Rambault, Doz-Deblauwe et al. 2021). This apparent discrepancy can be explained by the different methods used in the two studies. In our previous work we performed the phagocytosis assay on PMNs purified by fluorescence activated cell sorting, a process triggering cell activation. By contrast, in the present study, we worked with total milk or blood cells to assess phagocytosis while avoiding PMN activation by cell sorting. This simpler and less harsh method for PMN purification allowed us to confirm that blood MHC-II^pos^ PMNs possessed higher phagocytosis power than MHC-II^neg^ PMNs. Moreover, we could also demonstrate by using an *E. coli* strain carrying the GFP reporter, that superior phagocytosis by MHC-II^pos^ PMNs as compared to MHC-II^neg^ was retained after recruitment to the mammary gland. Both *E*.*coli* P4-GFP strain and pHrodo *E*.*coli* bioparticules were used to measure PMNs phagocytosis activity and gave similar results (data not shown). We think these data are interesting since PMNs are the most important killer of bacteria invading the mammary gland during mastitis and early control of infection mainly relies on this population especially during infection with *E. coli* or *Staphylococcus aureus* which are widespread mastitis pathogens (Rainard and Riollet 2003). Therefore, a better understanding on the proportion, kinetics and level of recruitment as well as regulation of different PMNs subsets recruited to the mammary gland could help to implement better control measures for example by introducing such markers in genetic selection programs. More work is needed though and in the low number of animals that we analyzed in this study we could not correlate the levels of MHC-II^neg^ PMNs versus MHC-II^pos^ PMNs with differential control of bacteria multiplication in the mammary gland during the longitudinal study of clinical mastitis.

We observed that MHC-II^pos^ PMNs reached milk both during clinical and subclinical forms of mastitis. MHC-II^pos^ PMNs accumulated in high numbers in milk as compared to circulation in blood, particularly during subclinical mastitis. We cannot explain yet the reason for such finding. Interestingly, we observed during this work that animals with highest proportion of MHC-II^pos^ PMNs in milk were those with repetitive high milk SCC recorded over several weeks or months (data not shown), which could suggest that this subset was associated to chronicity of mastitis. However, this interpretation must be taken with caution. Milk control is routinely applied in animal facilities or farms and abnormal SCC appears as one parameter measured of the whole mammary gland leading to exclusion of the animal until recovery. Thus, since we did not have the information about SCC per quarter for these animals, we could not accurately conclude that MHC-II^pos^ PMNs numbers in each quarter analyzed were correlated to mastitis caused by the same pathogen in the same quarter. The highly significant correlation of MHC-II^pos^ PMNs with T lymphocytes in milk supports the hypothesis that MHC-II^pos^ PMNs are involved in chronic mastitis and may be involved in the control of the adaptive immune response in the long-term. Also, we have recently demonstrated the capacity of blood MHC-II^pos^ PMNs to suppress T cell proliferation (Rambault, Doz-Deblauwe et al. 2021). Therefore, although it is not possible to state that MHC-II^pos^ PMNs are associated to chronic infection, this hypothesis remains interesting. Unfortunately, due to technical reasons, we could not demonstrate that milk MHC-II^pos^ PMNs suppressed T cells’ proliferation. We attempted to sort MHC-II^pos^ PMNs from milk both by flow cytometry and incubation with magnetic beads (Rambault, Borkute et al. 2021) but the high activation state and poor viability status of the cells after such procedures did not lead to conclusive data.

Several studies have reported that human PMNs expressing MHC-II behave as antigen-presenting cells (Culshaw, Millington et al. 2008, Abi Abdallah, Egan et al. 2011). Blood PMNs expressing MHC-II have also been reported in human patients with chronic inflammatory disease, but not during acute bacterial infections (Iking-Konert, Wagner et al. 2002). Altogether, this suggests that MHC-II^pos^ are involved in chronic mastitis and/or modeling of the adaptive immune phase of the immune response by interacting with T cells.

Nowadays, differential leukocytes count is investigated as a new tool for mastitis diagnosis (Schwarz, Diesterbeck et al. 2011, Damm, Holm et al. 2017, Goncalves, Lyman et al. 2017). Proportion of leukocytes populations provides more precise information about the inflammatory status of the mammary gland, but the description of cell types is limited to total PMNs, macrophages and lymphocytes with no information about subsets (Pilla, Schwarz et al. 2012, Godden, Royster et al. 2017, Kirkeby, Toft et al. 2020). A better understanding of the involvement of MHC-II^pos^ neutrophils in mastitis would improve differential counts and pave the way for identification of potential biomarkers of chronic mastitis.

To conclude, we believe that the diversity of phenotypes and functions of PMNs needs to be investigated more and we advocate that the classical description of PMNs as a homogeneous cell type in cattle must be revisited. Mastitis is still one of the most important issue for dairy farming; thus, investigating how MHC-II^pos^ PMNs behave during chronical mastitis would bring new views on the physiopathology of this costly disease. Our study also suggests that new tools such as differential counting used for mastitis diagnosis can be improved with a more accurate characterization of PMNs subsets. This could also open the way to the discovery of new biomarkers of chronic inflammation.

## Acknowledgments

This work was supported by the EGER program of APIS-GENE (MASTICELLS). MR conducts her doctoral project under a CIFRE agreement (industrial agreement of learning/training by research, CIFRE N°2019/0776) signed with Institut de l’Elevage IDELE. We warmly thank people working in the farms and experimental units involved in this project for milk and blood sampling and animal care, Manon Gillier and the “milk team” (Ferme experimentale des Trinottières, Montreuil-sur-Loir, France), Fréderic Launay and Sarah Barbey (Domaine expérimental du Pin-au-Haras), Eric Briant and his team (UE-PAO, INRAE, Nouzilly). We also thank Mickaël Riou and Noémie Perrot (PFIE, INRAE, Nouzilly) for complete blood count, Yves Le Vern and Alix Sausset for milk PMNs sorting assays. Plasmid pFPV25.1 was obtained from Dr Rodrigo Guabiraba (INRAE, Nouzilly). The authors declare that the research was conducted in the absence of any commercial or financial relationships that could be construed as a potential conflict of interest.

